# Location of phosphorylation sites within long polypeptide chains by binder-assisted nanopore detection

**DOI:** 10.1101/2024.04.29.590540

**Authors:** Wei-Hsuan Lan, Hanxiao He, Hagan Bayley, Yujia Qing

**Affiliations:** Department of Chemistry, University of Oxford, Oxford, UK

## Abstract

The detection and mapping of protein phosphorylation sites are essential for understanding the mechanisms of various cellular processes and for identifying targets for drug development. The study of biopolymers at the single-molecule level has been revolutionized by nanopore technology. In this study, we detect protein phosphorylation within long polypeptides (>600 amino acids), after the attachment of binders that interact with phosphate monoesters; electro-osmosis is used to drive the tagged chains through engineered protein nanopores. By monitoring the ionic current carried by a nanopore, phosphorylation sites are located within individual polypeptide chains, providing a valuable step toward nanopore proteomics.

## Introduction

Post-translational modifications (PTMs) of proteins are pivotal in cell regulation and typically involve the enzymatic addition of chemical groups to amino acid side chains^1^. Phosphorylation, the process of adding a phosphate group to predominantly serine, threonine, and tyrosine residues, is the most prevalent PTM, with over ∼10^6^ phosphorylation sites that account for >60% of all reported PTMs^1^. Dysregulation of phosphorylation is commonly associated with diseases, such as cancer, Parkinson’s, and Alzheimer’s^2^. For example, tau proteins in pathological lesions of Alzheimer’s are heterogeneously and highly phosphorylated, with more than 50 identified phosphorylation sites^3^. Bottom-up mass spectrometry is routinely applied to detect PTMs on peptide fragments derived from disease-related proteins, but faces challenges to determine if widely separated modifications, whether identical or distinct, are present on the same polypeptide chain. For example, the cross-talk between phosphorylation and O-GlcNAcylation was reported to regulate subcellular localization of proteins, such as tau^4^. However, there lacks a straightforward technique to correlate their presence at distant sites at the single-protein level^5^. Nanopore nucleic acid sequencing has emerged as a powerful technology to provide ultra-long DNA or RNA reads for long-range correlation of genomic or transcriptomic features^6,7^. Single-molecule sensing using protein nanopores therefore holds great potential for single-molecule analysis of full-length proteoforms^8–12^. Electro-osmosis has been demonstrated to propel unfolded polypeptides through nanopores^13–15^ and PTMs deep within long polypeptide chains have been located during translocation^14^. This work is a first step towards the label-free analysis of modified proteins extracted from biological samples^14^. In parallel, the identification of PTMs on short peptides (up to ∼30 amino acids) has been achieved^16–19^, either when the peptides are sensed as a whole or when a peptide is transported through the pore as a conjugate to an oligonucleotide^18^. Although PTMs containing branched structures (e.g. glycans) or entire proteins (e.g. ubiquitin) might be challenging to detect on polypeptides translocating through nanopores of ∼1 to 2 nm in internal diameter, >80% of the ∼400 PTM types are small (<∼300 Da) or narrow in shape^1^. In addition, nanopores with wider internal geometries (e.g ClyA) might be applicable for sensing bulkier PTMs. To demonstrate the broad applicability of the approach, we previously detected three PTMs (phosphorylation, glutathionylation, and glycosylation) on full-length proteins when segments of singly modified thioredoxin (Trx)-linker concatemers were stalled during translocation through a nanopore^14^. To our surprise, the glutathionylation and phosphorylation, placed at sites two amino acids apart, produced similar current blockades and noise patterns^14^. This prompts the question on how many PTMs can be discriminated among the 400 different natural PTMs identified so far by their perturbation to the ionic current driven through a protein nanopore^1^. To distinguish PTMs of similar electrical signatures or to allow targeted detection of specific PTMs, we seek to use PTM-specific binders to generate distinct current characteristics. To this end, we have explored a phosphorylation-specific reversible chemical binder, Phos-tag, which binds selectively and strongly to phosphate monoesters when complexed with zinc ions (e.g., for phosphoserine or phosphotheronine residues within model peptides, *K*_d_ = ∼0.7 µM; for phosphotyrosine residues within model peptides, 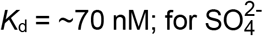, *K*_d_ = ∼130 µM; for Cl^-^, *K*_d_ = ∼2 mM)^20,21^. Phos-tag produced distinctive modulation of the associated ionic current as phosphorylated polypeptide chains were translocated through an engineered nanopore, allowing the location of phosphorylation sites within long polypeptide chains.

## Results and Discussion

In our previous research, we employed an anion-selective α-hemolysin (αHL) mutant (NN-113R)_7_ (permeability ratio 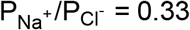)^22^ to generate electro-osmotic flow, thereby driving the capture, linearization, and translocation of polypeptide chains. We identified and located PTMs on long polypeptide chains of up to nine thioredoxin units (Trx, 108 amino acids (aa)) connected by linkers (29 aa)^14^. Each Trx units within the Trx-linker concatemers had the two catalytic cysteines removed (Trx: C32S/C35S)^8^. Chaotropic reagents (e.g. guanidinium chloride, GdnHCl, or urea) at non-denaturing concentrations were used to promote the co-translocational unfolding^14^. During the electro-osmotic translocation of the Trx-linker concatemers, features comprising three levels were seen (A1, A2 and A3) (**Figure 1a, b**). We provisionally assigned level A1 to be produced by the nanopore containing a threaded linker ahead of a folded Trx unit, level A2 to be produced when a partly unfolded C-terminus of a Trx unit extended into the nanopore, and level A3 to be produced by the spontaneous unfolding and passage of the remaining Trx polypeptide chain through the nanopore. In the presence of a PTM in the linker, a phosphate group (P) for instance, level A1 exhibited a smaller percentage residual current (*I*_res%_) value and higher root-mean-square noise (*I*_r.m.s.)_ ^14^ (**Figure 1b**).

**Figure 1.**
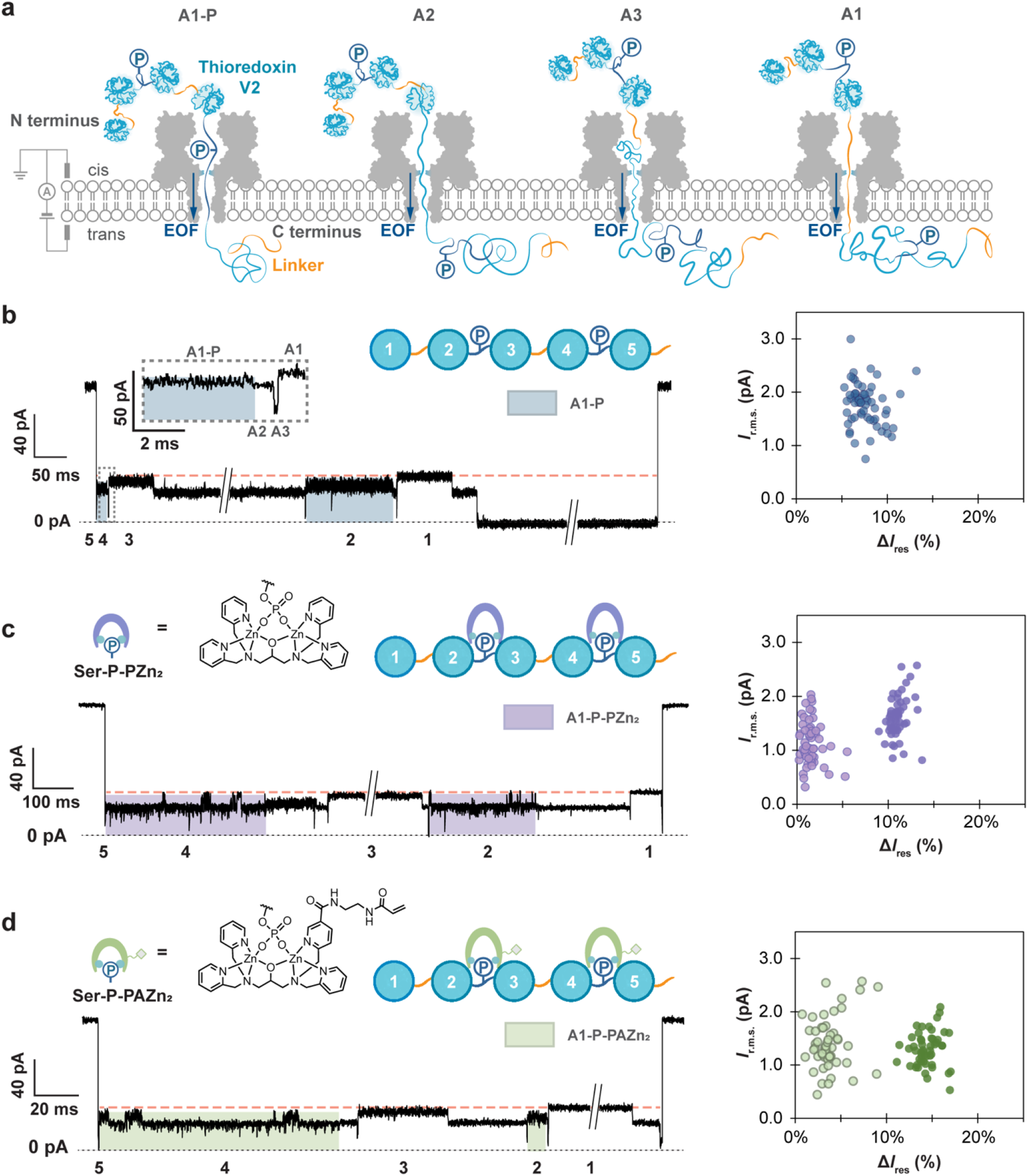
Detection of serine phosphate bound to Phos-tag in a polypeptide chain. **a**, Monitoring the Trx-linker pentamer traversing the α-hemolysin nanopore (NN-113R)_7_. The Trx-linker pentamer contained two RRAS sequences within the second and fourth linkers, which were phosphorylated on serine. **b**, Left: Phosphorylated serine residues (Ser-P) 274 aa apart on a Trx-linker pentamer were detected. Level A1 for the linker between Trx unit 3 and unit 4 showed a slightly lower *I*_res%_ compared to unmodified segments, such as the linker between first and second Trx (average *I*_res%_ of unmodified segments is shown in orange dash). This difference was attributed to the additional amino acid sequence in the third linker (**Table S1**). Right: Scatter plots of *I*_r.m.s._ and Δ*I*_res%_ for individual translocation events, Δ*I*_res%_ = <*I*_res%_(A1, Trx-linker)> – *I*_res%_(A1-P), where <*I*_res%_(A1, Trx-linker)> is the mean *I*_res%_ value of the remaining A1 levels for unmodified repeat units within an individual translocation event. If there were two Ser-P detected in different segments within a single translocation event, they were analyzed individually. **c**, Left: Phos-tag dizinc complexes bound to phosphoserine generated alternating current levels (A1-P-PZn_2_). Right: Scatter plots of *I*_r.m.s._ and Δ*I*_res%_ for individual translocation events. Data points in light purple are the *I*_r.m.s._ and Δ*I*_res%_ values for the higher level of the two-level A1 state (A1-P-PZn_2_-H), while data points in dark purple are the *I*_r.m.s._ and Δ*I*_res%_ values for the lower level of the two-level A1 state (A1-P-PZn_2_-L). **d**, Left: Phos-tag-acrylamide dizinc complexes bound to serine phosphate produced alternating current levels (A1-P-PAZn_2_). Right: Scatter plots of *I*_r.m.s._ and Δ*I*_res%_ for individual translocation events. Data points in light green are the *I*_r.m.s._ and Δ*I*_res%_ values for the higher level of the two-level A1 state (A1-P-PAZn_2_-H), while data points in dark green are the *I*_r.m.s._ and Δ*I*_res%_ values for the lower level of the two-level A1 state (A1-P-PAZn_2_-L). If there were two A1-P-PZn_2_ or A1-P-PAZn_2_ detected in different segments in a single translocation event, they were analyzed individually. Conditions in **b**: 10 mM HEPES, pH 7.2, 750 mM GdnHCl, 2.37 μM Trx-linker pentamer (cis). Conditions in **c**: 10 mM HEPES, pH 7.2, 750 mM GdnHCl, 2.37 μM Trx-linker pentamer (cis), 118.5 μM Phos-tag (cis), 237 μM ZnCl_2_ (cis). Condition in **d**:10 mM HEPES, pH 7.2, 750 mM GdnHCl, 2.37 μM Trx-linker pentamer (cis), 118.5 μM Phos-tag-acrylamide (cis), 237 μM ZnCl_2_ (cis). All the measurement were conducted at +140 mV (trans) and 23 ± 1 °C.

Here, we have examined the detection of phosphorylation in association with phosphate-specific binders: Phos-tag dizinc complex (PZn_2_) and Phos-tag-acrylamide dizinc complex (PAZn_2_). Phos-tag is commonly immobilized in SDS-PAGE gels to detect phosphoproteins^23,24^ and applied to generate mass shifts in MALDI-TOF mass spectrometry^25^. We constructed a Trx-linker pentamer containing two phosphorylation sites (RRAS) in the second and fourth linkers (**Figure 1a, Table S1 and Figure S1**), which were phosphorylated on serine by the catalytic subunit of protein kinase A (**Figure S2**). Phosphorylated polypeptides were captured, unfolded, and translocated by electro-osmosis through the (NN-113R)_7_ αHL pore. GdnHCl (750 mM) was employed to accelerate the co-translocational unfolding. Consistent with prior findings, translocation of the pentamer, C-terminus first, generated current patterns with a maximum of 4 A1-A3 repeats following an initial spike (**Figure 1b**). The spike to around 0 pA at the beginning of nearly all the translocation events was attributed to rapid unfolding and translocation of the first C-terminal Trx-linker unit. While only ∼6% of the doubly phosphorylated Trx-linker pentamers produced 5 repeating A1-A3 features, >72% of the recorded translocation events contained at least one A1 level with a reduced *I*_res%_ value and a higher *I*_r.m.s._, compared to A1 levels for unmodified segments (**Table S2**). These characteristics aligned with the electrical profiles previously identified for a phosphorylated linker and therefore assigned as level A1-P. In events where 5 repeats of A1-A3 features were observed, the level A1-P was recorded for both the second and fourth units, consistent with the presence of two phosphorylated serine residues (Ser-P) within the second and fourth linkers, 274 amino acids apart within the polypeptide chain.

To determine whether the binding of PZn_2_ and PAZn_2_ to phosphates in the polypeptide chains could be identified during translocation, we pre-formed complexes of phosphorylated Trx-linker pentamer with PZn_2_ and PAZn_2_, individually, with a molar ratio of Trx-linker:Phos-tag or Phos-tag-acrylamide:ZnCl_2_ = 1:50:100 (10 mM HEPES, pH 7.2, 750 mM GdnHCl, 2.37 μM Trx-linker pentamer, 118.5 μM Phos-tag or Phos-tag-acrylamide, 237 μM ZnCl_2_), and added the mixture to the cis compartment of the recording chamber. While the unmodified segments exhibited A1 levels characteristic of the unphosphorylated linkers, the phosphorylated linkers generated a distinctive A1 state with an ionic current that alternated between two levels (**Figure 1c, d**). The percentage residual currents of the two levels resulted from PZn_2_ binding were both smaller compared to those from PAZn_2_ (Δ*I*_res%_ for the higher level of the alternating current steps = 1.6 ± 1.0% for PZn_2_ and 3.8 ± 1.7% for PAZn_2_; Δ*I*_res%_ for the lower level of the alternating steps = 11 ± 1% for PZn_2_ and 14 ± 1% for PAZn_2_) (**Figure 1c, d and Table S2)**. We attributed the larger current blockade to the acrylamide appendix in PAZn_2_ and continued to investigate the binder-assisted PTM detection using PAZn_2_. To verify whether the distinctive current feature stemmed from the association of PAZn_2_ and Ser-P in the Trx-linker pentamer, a competition assay was performed in which excess phosphoserine was introduced to compete for binding with PAZn_2_ (**Methods and Figure S3**). Nanopore characterization of the phosphorylated Trx-linker pentamers in complex with PAZn_2_ (preformed at a molar ratio of Trx-linker:Phos-tag-acrylamide:ZnCl_2_ = 1:50:100 (2.37 μM Trx-linker pentamer, 118.5 μM Phos-tag-acrylamide, 237 μM ZnCl_2_)) was first recorded for approximately 10 minutes (**Methods**). Subsequently, excess phosphoserine (237 μM) was added to the cis compartment, and another 10-minute recording was performed. Prior to the addition of phosphoserine, 81% of the A1-P levels (N = 29) exhibited two alternating steps. The frequency of these events dropped to 17% (N = 24) after the addition of phosphoserine. The current feature of alternating levels was not observed in the presence of surplus ZnCl_2_ without phos-tag-acrylamide (10 mM HEPES, pH 7.2, 750 mM GdnHCl, 2.37 μM Trx-linker pentamer, 2 mM ZnCl_2_) (**Figure S4**). These results suggest that state A1 with two interconverting levels arose from the binding of PAZn_2_ to Ser-P (henceforth A1-P-PAZn_2_). Transitions between a A1-P-PAZn_2_ level and a level with an ionic current closely similar to level A1-P were also detected (**Figure S5)**, which was attributed to the dissociation of PAZn_2_ from Ser-P while the phosphorylated polypeptide segment was within the pore. The two current levels in A1-P-PAZn_2_ likely reflect the two-step chelation of a phosphate monoester with PAZn_2_ ^26–28^. A kinetics analysis revealed that the level with larger current blockades (A1-P-PAZn_2_-L) had a mean dwell time that was ∼4 times longer than the level with smaller current blockades (A1-P-PAZn_2_-H) (<τ_A1-P-PAZn2-L_> = 11.6 ± 0.3 ms, <τ_A1-P-PAZn2-H_> = 3.3 ± 0.1 ms), indicating that level A1-P-PAZn_2_-L was the more stable binding state (**Table S3**). We suggest that level A1-P-PAZn_2_-L represents PAZn_2_ with both zinc ions chelated by phosphate oxygen atoms, and level A1-P-PAZn_2_-H, PAZn_2_ with only one zinc ion chelated by a phosphate oxygen atom.

To ensure complete detection of phosphorylated sites within Trx-linker pentamer, we further optimized the recording conditions using PAZn_2_. Increasing the binder concentration to 1000 equivalents over Trx-linker pentamer resulted in almost complete detection of PAZn_2_-bound Ser-P (83% at 10 mM HEPES, pH 7.2, 750 mM GdnHCl, 2.37 μM Trx-linker pentamer, 118.5 μM Phos-tag-acrylamide, 237 μM ZnCl_2_ (Trx-linker:Phos-tag-acrylamide:ZnCl_2_ = 1:50:100) versus 95% at 10 mM HEPES, pH 7.2, 750 mM GdnHCl, 2.37 μM Trx-linker pentamer, 2.37 mM Phos-tag-acrylamide, 4.74 mM ZnCl_2_ (Trx-linker:Phos-tag-acrylamide:ZnCl_2_ = 1:1000:2000)) (**Figure S6**). Reducing the cationic electrolyte, Gdn^+^, and the anionic buffering reagent, HEPES, resulted in complete converage (>99% at 2 mM HEPES, pH 7.2, 750 mM KCl, 2.37 μM Trx-linker pentamer, 237 μM Phos-tag-acrylamide, 474 μM ZnCl_2_ (Trx-linker:Phos-tag-acrylamide:ZnCl_2_ = 1:100:200) (**Figure S6**).

Next, we sought to determine if PAZn_2_ would enable us to distinguish phosphorylation from a PTM that exhibits a similar ionic blockade^14^. We constructed a Trx-linker pentamer with distinct modification sites in the second (RRASAA) and fourth (RRAAAC) linkers. We carried out phosphorylation and glutathionylation reactions sequentially to obtain a Trx-linker pentamer with Ser-P in the second linker and glutathionylated cysteine (Cys-GS) in the fourth linker (**Figure 2a**). In line with the characteristic current patterns recorded separately with Trx-linker nonamers containing a single Ser-P or Cys-GS residues within the same linker sequence^14^, the signals from Ser-P and Cys-GS within the same Trx-linker pentamer exhibited indistinguishable residual currents and noise when the second and fourth linkers were located within the pore (**Figure 2a**). Pleasingly, the introduction of PAZn_2_ altered the signal derived from the second linker to give a pattern similar to level A1-P-PAZn_2_, while the signal from the fourth linker was unchanged, allowing clear differentiation between phosphorylation and glutathionylation (**Figure 2b**).

**Figure 2.**
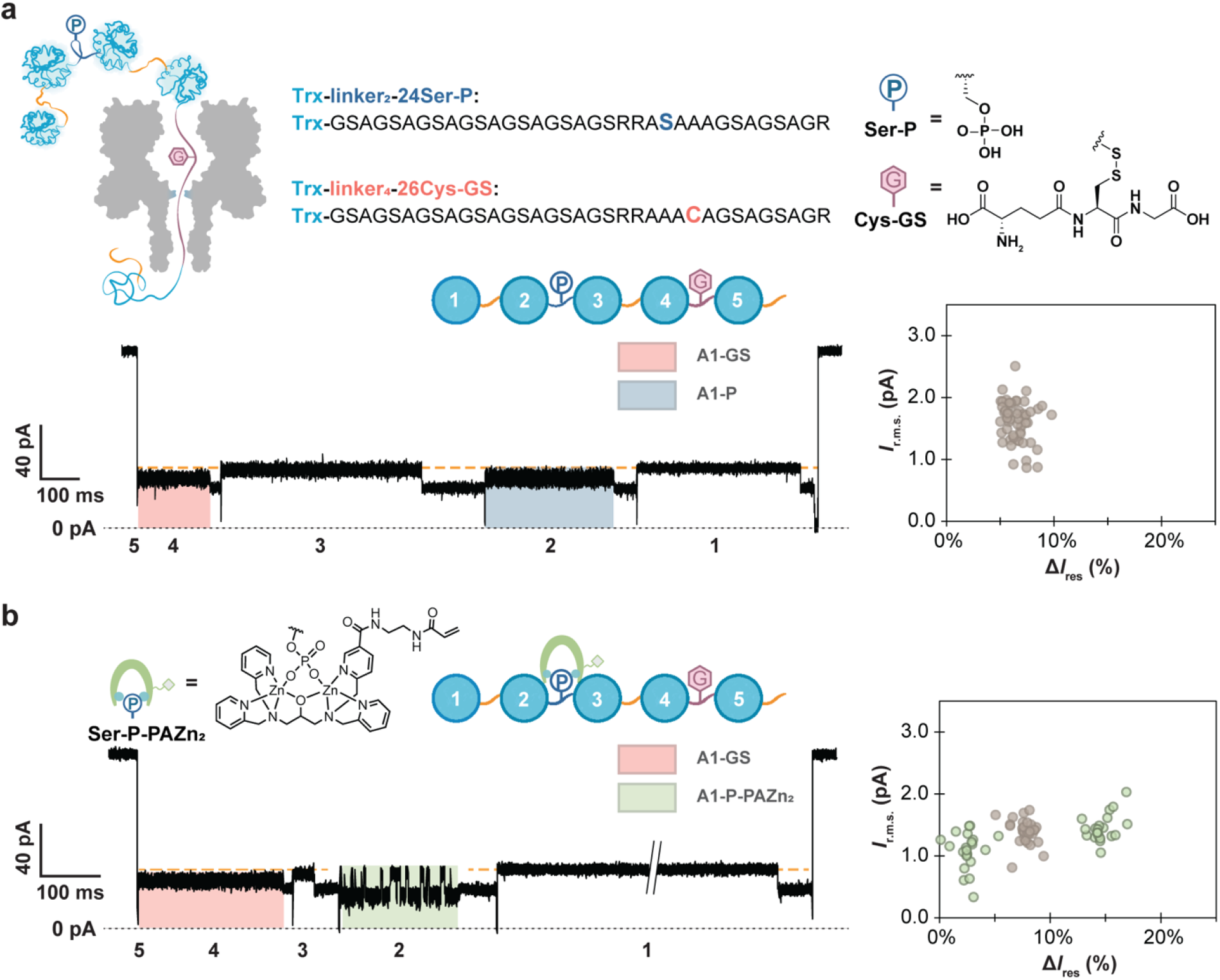
Detection of phosphorylation and glutathionylation in a Trx-linker pentamer in the presence of Phos-tag. **a**, The blockades and noises from Ser-P and Cys-GS cannot be readily discriminated. Top and bottom-left: Monitoring the phosphorylated and glutathionylated Trx-linker pentamer during translocation through a (NN-113R)_7_ αHL nanopore. The pentamer is phosphorylated on Ser-24 (Ser-P) of the second linker and glutathionylated on the Cys-26 (Cys-GS) of the fourth linker. Bottom-right: Scatter plots of *I*_r.m.s._ and Δ*I*_res%_ for individual translocation events, Δ*I*_res%_ = <*I*_res%_(A1, Trx-linker)> – *I*_res%_(A1*), where <*I*_res%_(A1, Trx-linker)> is the mean *I*_res%_ value of the remaining A1 levels for unmodified repeat units within an individual translocation event and *I*_res%_(A1*) is the *I*_res%_ value of A1-GS and A1-P. If there were two A1* detected in different segments within a single translocation event, they were analyzed individually. **b**, PAZn_2_ produced an additional current feature when bound to Ser-P. Left: Monitoring the phosphorylated and glutathionylated Trx-linker pentamer in the presence of PAZn_2_ during translocation through a (NN-113R)_7_ αHL nanopore. Right: Scatter plots of *I*_r.m.s._ and Δ*I*_res%_ for individual translocation events. Data points in gray are the *I*_r.m.s._ and Δ*I*_res%_ values for A1-GS and A1-P, while data points in green are the *I*_r.m.s._ and Δ*I*_res%_ values for the higher and lower level of the two-level A1 state (A1-P-PZn_2_-H and A1-P-PZn_2_-L). Conditions in **a**: 10 mM HEPES, pH 7.2, 750 mM GdnHCl, 2.37 μM Trx-linker pentamer (cis), +140 mV (trans), 23 ± 1 °C. Conditions in **b**: 10 mM HEPES, pH 7.2, 750 mM GdnHCl, 2.37 μM Trx-linker pentamer (cis), 118.5 μM Phos-tag-acrylamide (cis), 237 μM ZnCl_2_ (cis), +140 mV (trans), 23 ± 1 °C.

## Conclusion

Here, we demonstrate the nanopore detection of widely separated phosphorylation sites (e.g. >250 aa apart) within a polypeptide chain by using Phos-tag dizinc complex (PZn_2_) and Phos-tag-acrylamide dizinc complex (PAZn_2_). The binder created a distinct two-level current feature when phosphorylated polypeptide segments were inside the nanopore, which resembled current patterns observed during divalent cation chelation within an engineered αHL pore^26^ or with amino acids interacting with immobilized Ni^2+^ in an engineered nanopore^28^. We have saturated Ser-P with PAZn_2_ under the condition with excess PAZn_2_ (10 mM HEPES, pH 7.2, 750 mM GdnHCl, 2.37 μM Trx-linker pentamer, 2.37 mM Phos-tag-acrylamide, 4.74 mM ZnCl_2_ (Trx-linker:Phos-tag-acrylamide:ZnCl_2_ = 1:1000:2000)) in this work. The phosphorylation-specific current feature enabled the discrimination of phosphorylation from PTMs that produced similar current blockades, such as glutathionylation. We envision that combinations of PTM-specific binders will allow the simultaneous detection of multiple PTMs. Suitable binders should recognize PTMs irrespective of adjacent amino acid sequences, exhibit fast association and slow dissociation kinetics, and ideally generate characteristic current signatures, such as the subconductance states seen in this work, which facilitate the discrimination of PTMs. Given that most of PTMs are enzymatically installed and regulated, they tend to be located within the flexible or exposed regions of proteins^29^. For the small fraction of PTMs that are non-enzymatically installed within the buried regions of proteins (e.g. disulfide bonds), partial unfolding might occur in the presence of chaotropic reagents to allow binder association. So far, we have identified PTMs in polypeptide segments while they are transiently arrested within a nanopore. In our ongoing efforts to study biologically relevant protein targets, the use of bulky binders (e.g., antibodies) holds promise for temporarily halting protein translocation at the pore entrance, thereby mediating PTM identification in any region of a protein.

## Acknowledgements

This research was supported by the Wellcome Leap Delta Tissue Program, Oxford Nanopore Technologies. W.-H.L. is funded by an Oxford-Taiwan Graduate Studentship in partnership with a Department of Chemistry Scholarship, University of Oxford. For the purpose of open access, the authors have applied a CC BY public copyright licence to any Author Accepted Manuscript version arising from this submission.

## Author Contributions

W.-H.L., H.B., and Y.Q. designed the study; W.-H.L. and H.H. performed the experiments; and W.-H.L., H.B., and Y.Q. wrote the manuscript.

## Supporting Information

Tables S1–S3 and Figures S1–S6; experimental procedures and characterization data.

