## Supplemental information for "Location of phosphorylation sites within long polypeptide chains by binder-assisted nanopore detection"

### Table of Contents

**Table S1** Amino acid sequences of the thioredoxin-linker pentamers **3**

**Table S2** Percentage residual current ($I_{res\%}$) and root-mean-square noise ($I_{r.m.s.}$) characteristics of A1-P, A1-P-PZn$_2$, and A1-P-PAZn$_2$ **4**

**Table S3** Mean dwell times ($<\tau>$) derived by QuB for two-level A1-P-PAZn$_2$ **5**

**Figure S1** An SDS-polyacrylamide gel of the Trx-linker pentamer **6**

**Figure S2** ESI LC-MS characterization of Trx-linker pentamers **7**

**Figure S3** Fractions of phosphorylated linkers detected in the PAZn$_2$-bound state in the absence and presence of competing phosphoserine **8**

**Figure S4** An ionic current trace showing phosphorylated linkers in excess $Zn^{2+}$ generated similar A1-P current signals **9**

**Figure S5** A current trace showing transition between level A1-P-PAZn$_2$ and level A1-P when a phosphorylated segment was inside the (NN-113R)$_7$ nanopore **10**

**Figure S6** Fractions of phosphorylated linkers detected in the PAZn$_2$-bound state **11**

**Methods** **12**

**References** **15**

**Table S1. Amino acid sequences of the thioredoxin-linker pentamers**

| **(Trx-linker)$_{1,3,5}$(Trx-linker-24S26C)$_{2,4}$** | **Figure 1** |
|---|---|
| SDKIIHLTDDSFDTDVLKADGAILVDFWAEWSGPSKMIAPILDEIADEYQGKLTVAKLNIDQNPGT<br>APKYGIRGIPTLLLFKNGEVAATKVGALSKGQLKEFLDANLAGSAGSAGSAGSAGSAGSAGSAG<br>SAGSAGRSDKIIHLTDDSFDTDVLKADGAILVDFWAEWSGPSKMIAPILDEIADEYQGKLTVAKLN<br>IDQNPGTAPKYGIRGIPTLLLFKNGEVAATKVGALSKGQLKEFLDANLAGSAGSAGSAGSAGSA<br>GSAGS**RRASAC**AGSAGSAGRSDKIIHLTDDSFDTDVLKADGAILVDFWAEWSGPSKMIAPILDEI<br>ADEYQGKLTVAKLNIDQNPGTAPKYGIRGIPTLLLFKNGEVAATKVGALSKGQLKEFLDANLAGS<br>AGSAGSAGSAGSAGSAGSAGSAGSAGRSGTGGPRRRSDKIIHLTDDSFDTDVLKADGAILVDF<br>WAEWSGPSKMIAPILDEIADEYQGKLTVAKLNIDQNPGTAPKYGIRGIPTLLLFKNGEVAATKVG<br>ALSKGQLKEFLDANLAGSAGSAGSAGSAGSAGSAGS**RRASAC**AGSAGSAGRSDKIIHLTDDSF<br>DTDVLKADGAILVDFWAEWSGPSKMIAPILDEIADEYQGKLTVAKLNIDQNPGTAPKYGIRGIPTL<br>LLFKNGEVAATKVGALSKGQLKEFLDANLAGSAGSAGSAGSAGSAGSAGSAGSAGSAGRS | |
| **(Trx-linker)$_{1,3,5}$(Trx-linker-24S)$_2$(Trx-linker-26C)$_4$** | **Figure 2** |
| SDKIIHLTDDSFDTDVLKADGAILVDFWAEWSGPSKMIAPILDEIADEYQGKLTVAKLNIDQNPGT<br>APKYGIRGIPTLLLFKNGEVAATKVGALSKGQLKEFLDANLAGSAGSAGSAGSAGSAGSAGSAG<br>SAGSAGRSDKIIHLTDDSFDTDVLKADGAILVDFWAEWSGPSKMIAPILDEIADEYQGKLTVAKLN<br>IDQNPGTAPKYGIRGIPTLLLFKNGEVAATKVGALSKGQLKEFLDANLAGSAGSAGSAGSAGSA<br>GSAGS**RRASAA**AGSAGSAGRSDKIIHLTDDSFDTDVLKADGAILVDFWAEWSGPSKMIAPILDEI<br>ADEYQGKLTVAKLNIDQNPGTAPKYGIRGIPTLLLFKNGEVAATKVGALSKGQLKEFLDANLAGS<br>AGSAGSAGSAGSAGSAGSAGSAGSAGRSGTGGPRRRSDKIIHLTDDSFDTDVLKADGAILVDF<br>WAEWSGPSKMIAPILDEIADEYQGKLTVAKLNIDQNPGTAPKYGIRGIPTLLLFKNGEVAATKVG<br>ALSKGQLKEFLDANLAGSAGSAGSAGSAGSAGSAGS**RRAAAC**AGSAGSAGRSDKIIHLTDDSF<br>DTDVLKADGAILVDFWAEWSGPSKMIAPILDEIADEYQGKLTVAKLNIDQNPGTAPKYGIRGIPTL<br>LLFKNGEVAATKVGALSKGQLKEFLDANLAGSAGSAGSAGSAGSAGSAGSAGSAGSAGRS | |
| Blue: Trx; Yellow: linker; Orange: modified linker; **Pink**: sequence of the modification; White: restriction enzyme sites (KpnI and AvrII). | |

**Table S2. Percentage residual current ($I_{\text{res\%}}$) and root-mean-square noise ($I_{\text{r.m.s.}}$) characteristics of A1-P, A1-P-PZn$_2$, and A1-P-PAZn$_2$[a], [b]**

| | | $\Delta I_{\text{res\%}}$[a] | $I_{\text{r.m.s.}}$ / pA[b] | N |
|---|---|---|---|---|
| **A1-P** | | 7.6 ± 1.6% | 1.8 ± 0.4 | 53 concatemers<br>4 separate pores |
| **A1-P-PZn$_2$** | **A1-P-PZn$_2$-H** | 1.6 ± 1.0% | 1.1 ± 0.4 | 54 concatemers<br>6 separate pores |
| | **A1-P-PZn$_2$-L** | 11 ± 1% | 1.6 ± 0.4 | |
| **A1-P-PAZn$_2$** | **A1-P-PAZn$_2$-H** | 3.8 ± 1.7% | 1.4 ± 0.5 | 51 concatemers<br>5 separate pores |
| | **A1-P-PAZn$_2$-L** | 14 ± 1% | 1.3 ± 0.3 | |

**[a]** $\Delta I_{\text{res\%}} = <I_{\text{res\%}}(\text{A1, Trx-linker})> - I_{\text{res\%}}(\text{A1-P})$, $<I_{\text{res\%}}(\text{A1, Trx-linker})> - I_{\text{res\%}}(\text{A1-P-PZn}_2\text{-H})$, $<I_{\text{res\%}}(\text{A1, Trx-linker})> - I_{\text{res\%}}(\text{A1-P-PZn}_2\text{-L})$, $<I_{\text{res\%}}(\text{A1, Trx-linker})> - I_{\text{res\%}}(\text{A1-P-PAZn}_2\text{-H})$, or $<I_{\text{res\%}}(\text{A1, Trx-linker})> - I_{\text{res\%}}(\text{A1-P-PAZn}_2\text{-L})$. For a C terminus-first translocation event, $<I_{\text{res\%}}(\text{A1, Trx-linker})>$ was determined as the mean $I_{\text{res\%}}$ value of the unmodified A1 levels within an individual translocation event. $I_{\text{res\%}}$(A1-P) was determined for the A1 level of the modified linker and appeared once or twice per translocating pentamer. $I_{\text{res\%}}$(A1-P-PZn$_2$-H) and $I_{\text{res\%}}$(A1-P-PZn$_2$-L), as well as $I_{\text{res\%}}$(A1-P-PAZn$_2$-H) and $I_{\text{res\%}}$(A1-P-PAZn$_2$-L), were determined for the higher and lower levels of the two-level A1-P-PZn$_2$ and A1-P-PAZn$_2$ states, which occurred once or twice per translocating pentamer. If two A1-P, A1-P-PZn$_2$ or A1-P-PAZn$_2$ were detected in a single translocation event, they were analyzed individually. Conditions: 10 mM HEPES, pH 7.2, 750 mM GdnHCl, +140 mV (trans), 23 ± 1 °C.

**[b]** Root-mean-square noise values ($I_{\text{r.m.s.}}$) were measured from current traces after a post-recording filter of 2 kHz. $I_{\text{r.m.s.}}$ was normalised for the noise of each pore ($I_{\text{r.m.s.}}^2 = I_{\text{r.m.s.}}(\text{A1-P})^2 - I_{\text{r.m.s.}}(\text{open pore})^2$, $I_{\text{r.m.s.}}^2 = I_{\text{r.m.s.}}(\text{A1-P-PZn}_2\text{-H})^2 - I_{\text{r.m.s.}}(\text{open pore})^2$, $I_{\text{r.m.s.}}^2 = I_{\text{r.m.s.}}(\text{A1-P-PZn}_2\text{-L})^2 - I_{\text{r.m.s.}}(\text{open pore})^2$, $I_{\text{r.m.s.}}^2 = I_{\text{r.m.s.}}(\text{A1-P-PAZn}_2\text{-H})^2 - I_{\text{r.m.s.}}(\text{open pore})^2$, $I_{\text{r.m.s.}}^2 = I_{\text{r.m.s.}}(\text{A1-P-PAZn}_2\text{-L})^2 - I_{\text{r.m.s.}}(\text{open pore}^2)$).

**Table S3. Mean dwell times ($\langle\tau\rangle$) derived by QuB[a] for two-level A1-P-PA$Zn_2$[b]**

| Voltage (trans) | +140 mV | |
|---|---|---|
| **$\langle\tau_{A1\text{-}P\text{-}PAZn2\text{-}H}\rangle$ / ms** | 3.3 ± 0.1 | N = 224 |
| **$\langle\tau_{A1\text{-}P\text{-}PAZn2\text{-}L}\rangle$ / ms** | 11.6 ± 0.3 | N = 234 |

**[a]** Dwell time analysis was performed by using the maximum interval likelihood algorithm of QuB[1,2].

**[b]** Conditions: 10 mM HEPES, pH 7.2, 750 mM GdnHCl, 2.37 μM Trx-linker pentamer (cis), 118.5 μM Phos-tag-acrylamide (cis), 237 μM $ZnCl_2$ (cis), +140 mV (trans), 23 ± 1 °C. The C terminus-first translocations were recorded for Trx-linker pentamers through a single $(NN\text{-}113R)_7$ nanopore.

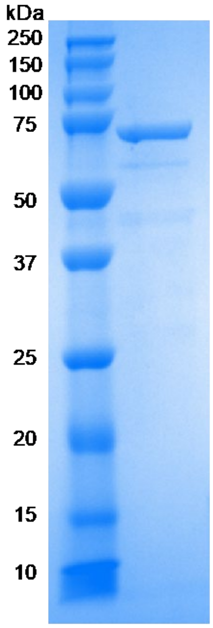


**Figure S1. An SDS-polyacrylamide gel of the Trx-linker pentamer**. (Trx-linker)$_{1,3,5}$(Trx-linker-24S26C)$_{2,4}$: 71 kDa.

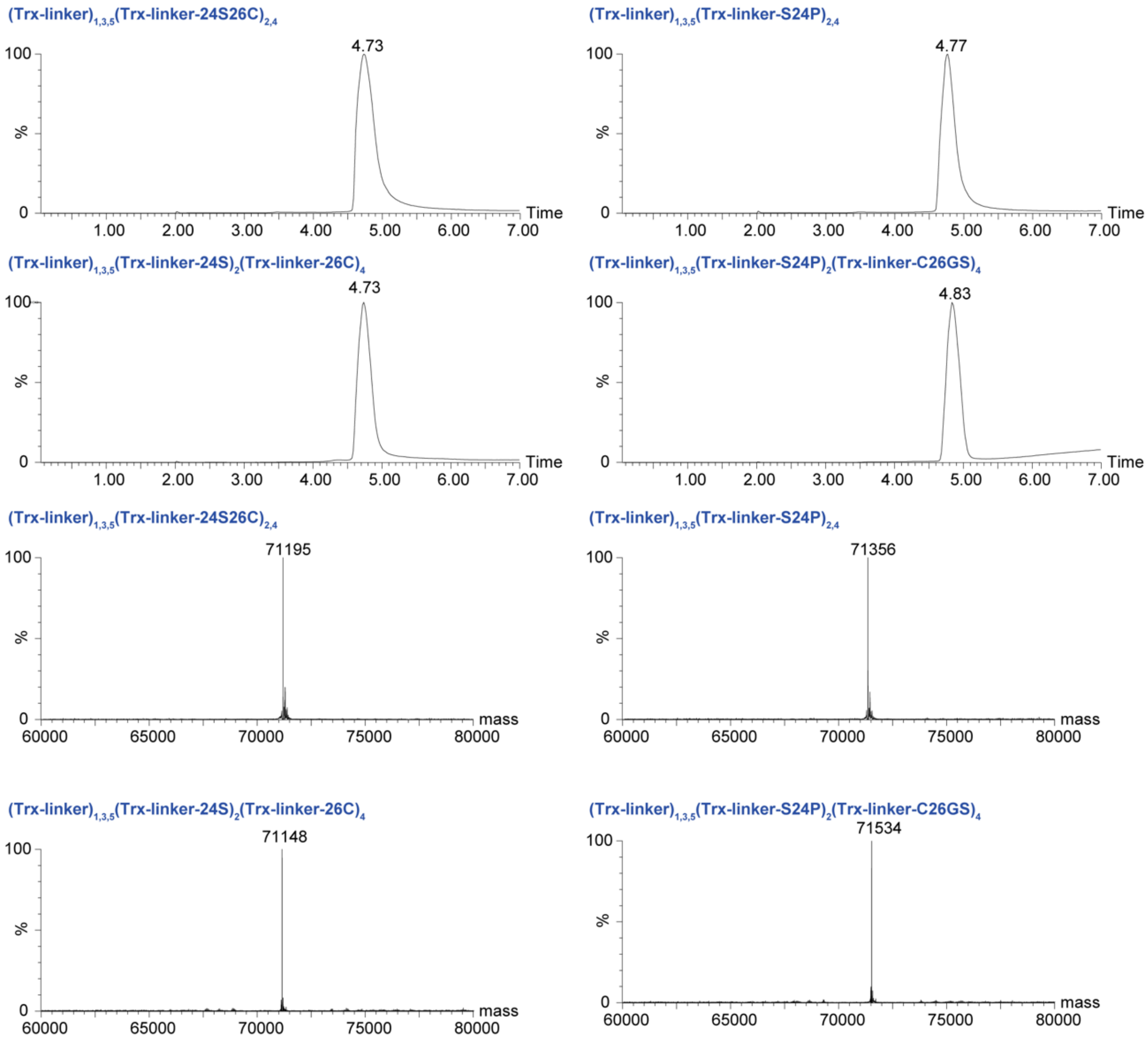


**Figure S2. ESI LC-MS characterization of Trx-linker pentamers**. LC-MS chromatograms (top) and deconvoluted ESI-MS spectra (bottom). (Trx-linker)$_{1,3,5}$(Trx-linker-24S26C)$_{2,4}$: mass = 71197 Da (calc) and 71195 Da (obs); (Trx-linker)$_{1,3,5}$(Trx-linker-S24P)$_{2,4}$: mass = 71356 Da (calc) and 71356 Da (obs); (Trx-linker)$_{1,3,5}$(Trx-linker-24S)$_2$(Trx-linker-26C)$_4$: mass = 71149 Da (calc) and 71148 Da (obs); (Trx-linker)$_{1,3,5}$(Trx-linker-S24P)$_2$(Trx-linker-C26GS)$_4$: mass = 71534 Da (calc) and 71534 Da (obs).

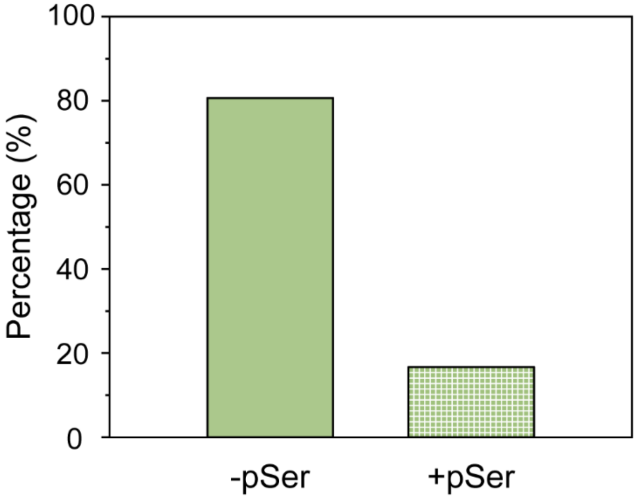


**Figure S3. Fractions of phosphorylated linkers detected in A1-P-PAZn$_2$ states in the absence and presence of competing phosphoserine.** Before pSer addition, 81% of phosphorylated linkers were detected in A1-P-PAZn$_2$ states (29 events). After pSer addition, 16% of phosphorylated linkers were detected in A1-P-PAZn$_2$ states (24 events).

Fractions (%) of phosphorylated linkers detected in A1-P-PAZn$_2$ states were calculated as:

Percentage =

$$\frac{\text{phosphorylated linkers detected in A1-P-PAZn}_2\text{ states}}{\text{phosphorylated linkers detected in A1-P-PAZn}_2\text{ states} + \text{phosphorylated linkers detected in A1-P states}}$$

If two phosphorylated linkers detected in A1-P or PAZn$_2$-bound states appeared in a single translocation event, they were analyzed individually. Conditions before adding pSer: 10 mM HEPES, pH 7.2, 750 mM GdnHCl, 2.37 μM Trx-linker pentamer (cis), 118.5 μM Phos-tag-acrylamide (cis), 237 μM $ZnCl_2$ (cis), +140 mV (trans), 23 ± 1 °C. Conditions after adding pSer: 10 mM HEPES, pH 7.2, 750 mM GdnHCl, 2.37 μM Trx-linker pentamer (cis), 118.5 μM Phos-tag-acrylamide (cis), 237 μM $ZnCl_2$ (cis), 237 μM pSer (cis), +140 mV (trans), 23 ± 1 °C.

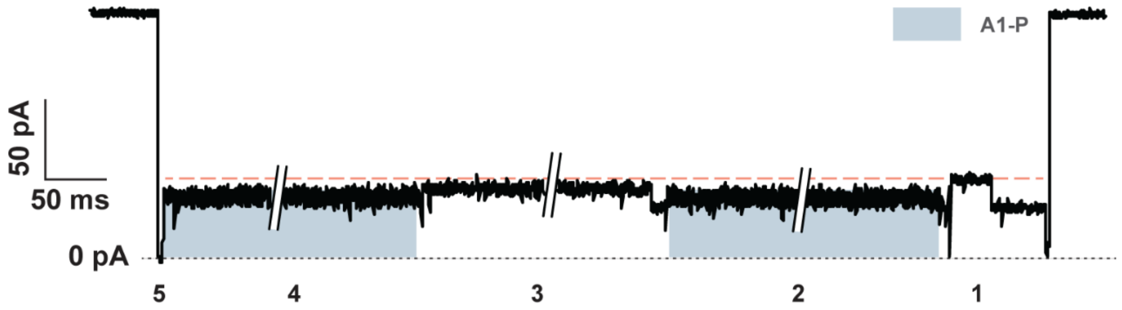


**Figure S4. An ionic current trace showing phosphorylated linkers in excess $Zn^{2+}$ generated similar A1-P current signals.** In the presence of excess $Zn^{2+}$ (~850 eq. over Trx-linker pentamer), phosphorylated linkers (the second and the fourth linkers) did not produce alternating levels during translocation. Conditions: 10 mM HEPES, pH 7.2, 750 mM GdnHCl, 2.37 μM Trx-linker pentamer (cis), 2 mM $ZnCl_2$ (cis), +140 mV (trans), 23 ± 1 °C.

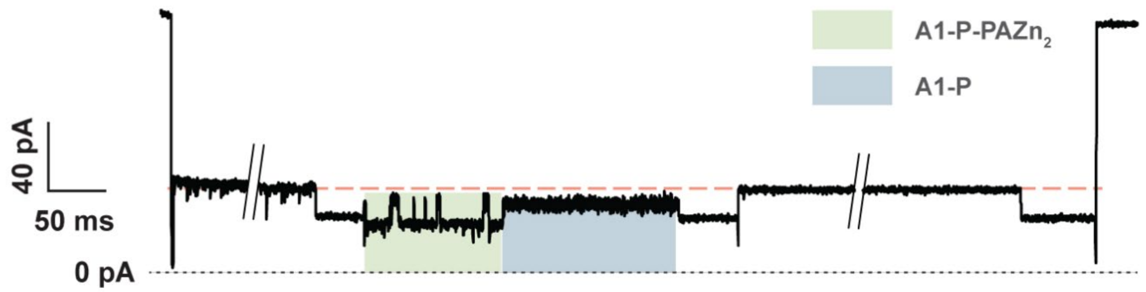


**Figure S5. A current trace showing transition between level A1-P-PAZn$_2$ and level A1-P when a phosphorylated segment was inside the (NN-113R)$_7$ nanopore.** Conditions: 10 mM HEPES, pH 7.2, 750 mM GdnHCl, 2.37 μM Trx-linker pentamer (cis), 118.5 μM Phos-tag-acrylamide (cis), 237 μM ZnCl$_2$ (cis), +140 mV (trans), 23 ± 1 °C.

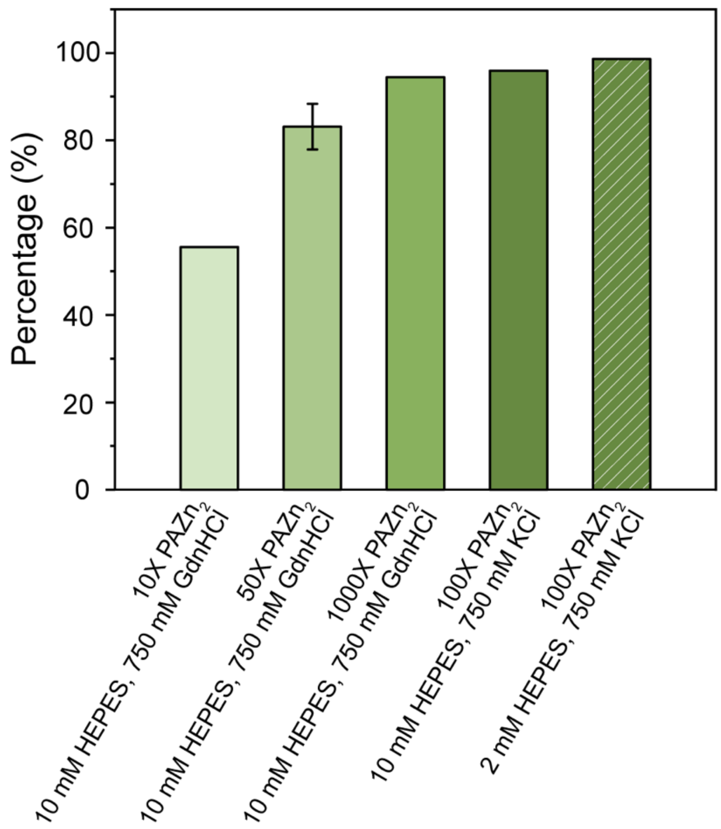


**Figure S6. Fractions of phosphorylated linkers detected in the $PAZn_2$-bound state.** The fractions of phosphorylated linkers detected in A1-P-$PAZn_2$ states were tested in five different conditions.

Fractions (%) of phosphorylated linkers detected in A1-P-$PAZn_2$ states were calculated as:

$$\text{Percentage} = \frac{\text{phosphorylated linkers detected in A1-P-PAZn}_2\text{ states}}{\text{phosphorylated linkers detected in A1-P-PAZn}_2\text{ states} + \text{phosphorylated linkers detected in A1-P states}}$$

If two phosphorylated linkers detected in A1-P or $PAZn_2$-bound states appeared in a single translocation event, they were analyzed individually. Conditions from left to right: 10 mM HEPES, pH 7.2, 750 mM GdnHCl, 2.37 μM Trx-linker pentamer (cis), 23.7 μM Phos-tag-acrylamide (cis), 47.4 μM $ZnCl_2$ (cis); 10 mM HEPES, pH 7.2, 750 mM GdnHCl, 2.37 μM Trx-linker pentamer (cis), 118.5 μM Phos-tag-acrylamide (cis), 237 μM $ZnCl_2$ (cis); 10 mM HEPES, pH 7.2, 750 mM GdnHCl, 2.37 μM Trx-linker pentamer (cis), 2.37 mM Phos-tag-acrylamide (cis), 4.74 mM $ZnCl_2$ (cis); 10 mM HEPES, pH 7.2, 750 mM KCl, 2.37 μM Trx-linkes pentamer (cis), 237 μM Phos-tag-acrylamide (cis), 474 μM $ZnCl_2$ (cis); 2 mM HEPES, pH 7.2, 750 mM KCl, 2.37 μM Trx-linker pentamer (cis), 237 μM Phos-tag-acrylamide (cis), 474 μM $ZnCl_2$ (cis). All the measurement were conducted at +140 mV (trans), 23 ± 1 °C.

#### Methods

##### Construction of His-SUMO-tagged Trx-linker pentamer genes

Reagents were purchased from NEB (New England Biolabs), unless otherwise stated. His-SUMO-tagged Trx-linker pentamer genes were prepared as previously described[3,4]. Two variants of His-SUMO-tagged Trx-linker pentamers were prepared to contain two phosphorylation sites within the second and fourth linkers (His-SUMO-tagged (Trx-linker)$_{1,3,5}$(Trx-linker-24S26C)$_{2,4}$) or one phosphorylation site within the second linker and one glutathionylation site within the fourth linker (His-SUMO-tagged (Trx-linker)$_{1,3,5}$(Trx-linker-24S)$_2$(Trx-linker-26C)$_4$).

##### Expression and purification of Trx-linker pentamers

Plasmids encoding the Trx-linker pentamer were transformed into BLR(DE3) competent cells (Novagen), which were cultivated in Luria broth (LB) supplemented with carbenicillin (100 µg/mL) at 37 °C with constant agitation at 250 rpm. Protein expression was induced in the exponential growth phase ($OD_{600}$ = 0.6 to 0.8) by adding isopropyl-β-D-1-thiogalactopyranoside (IPTG) to a final concentration of 0.5 mM. After 6 h, cells were harvested by centrifugation (at 5,000 g for 10 minutes), resuspended in a binding buffer (containing 30 mM Tris-HCl, 250 mM NaCl, 25 mM imidazole, pH 7.2) supplemented with a protease inhibitor cocktail (cOmplete™, EDTA-free, Roche), and lysed by sonication. Cell debris was removed by centrifugation at 20,000 g for 40 min, and the supernatant was loaded onto a column packed with HisPur Ni-NTA Agarose Resin (5 mL, ThermoFisher) equilibrated with binding buffer (25 mM Tris-HCl, pH 7.5, 500 mM NaCl, 25 mM imidazole) and the flow through was re-applied 5 times. After washing with binding buffer, the hexahistidine (His6)-tagged protein was eluted with 12 mL elution buffer (25 mM Tris-HCl, pH 7.5, 500 mM NaCl, 500 mM imidazole) and dialysed (Slide-A-Lyzer G2 Dialysis Cassette, 10,000 MWCO 30 mL, ThermoFisher) for 2 h against 4 L of dialysis buffer (50 mM Tris-HCl, pH 8.0, 150 mM NaCl, 2 mM 1,4-dithio-D-threitol (DTT)), with continuous stirring at 4 °C, to remove imidazole. Then, His6-tagged Ulp1 protease, prepared as previously described[4], was injected into the dialysis cassette at a 1: 200 molar concentration ratio with respect to the Trx-linker pentamer. Afterwards, the cassette was transferred to DTT-free dialysis buffer (50 mM Tris-HCl, pH 8.0, 150 mM NaCl) overnight for SUMO-tag cleavage. The cassette was then transferred to DTT-free dialysis buffer for an additional 4 h. The dialysed protein was loaded onto a column packed with 5 mL HisPur Ni-NTA Agarose Resin equilibrated with dialysis buffer and the flow through was re-applied to the column 5 more times. The final flow through containing the His6-SUMO-free protein was aliquoted and flash frozen for storage at -80 °C. The mass of the protein was confirmed by electrospray ionization liquid chromatography-mass spectrometry (ESI LCMS) (**Figure S2**).

##### Phosphorylation of Trx-linker pentamers

Trx-linker pentamers containing two phosphorylation sites within the second and fourth linkers or a single phosphorylation site within the second linker were phosphorylated by the catalytic subunit of the cAMP-dependent protein kinase (PKA) (NEB). The Trx-linker pentamers at a concentration of 1 mg/mL were incubated with 25,000 units of cAMP-dependent protein kinase (PKA) catalytic subunit (NEB), which phosphorylates the RRAS motif on serine. The buffer used contained 50 mM TrisHCl, pH 7.5,10 mM $MgCl_2$, 0.1 mM EDTA, 4 mM DTT, 0.01% Brij 35, and 2 mM ATP at 30 °C for 1 h. Then, the mixture was further supplemented with an additional 2 mM ATP and 2 mM DTT, followed by incubation at 30 °C for one more hour. The phosphorylated Trx-linker pentamers were purified and concentrated by using centrifugal filters (Vivaspin 2 centrifugal concentrators MWCO 50 kDa). They were then aliquoted and flash frozen for storage at -20 °C

(10 mM HEPES, pH 7.2, and 750 mM KCl). Phosphorylation of the Trx-linker pentamers was verified by LCMS (**Figure S2**).

**Modification of cysteine on Trx-linker pentamers**

Trx-linker pentamers containing a phosphorylation site within the second linker and a glutathionylation site within the fourth linker were first phosphorylated following the steps described in the above section. To subsequently glutathionylate the singly phosphorylated Trx-linker pentamers, they were treated with tris(2-carboxyethyl)phosphine (TCEP, Sigma-Aldrich) (100 eq.) at 32 °C for 2 h in protein storage buffer (50 mM TrisHCl, 250 mM NaCl, pH 8.0) and then desalted with PD MiniTrap G-25 columns (Cytiva). The reduced proteins were reacted with oxidized glutathione (100 eq.) (Sigma-Aldrich) at 32 °C overnight in protein storage buffer before desalting (PD MiniTrap G-25 columns). The glutathionylated proteins were aliquoted, flash frozen, and stored at -20 °C.

**Phosphoserine competition assay**

Phos-tag-acrylamide was purchased from FUJIFILM Wako Chemicals Europe GmbH. The phosphorylated Trx-linker pentamer was mixed with Phos-tag-acrylamide dizinc complex with a molar ratio of Trx-linker:Phos-tag-acrylamide:$ZnCl_2$ = 1:50:100 and kept at room temperature for 15 min. The mixture was then added to the cis compartment of the recording chamber (final concentrations in the cis compartment: 2.37 μM Trx-linker pentamers, 118.5 μM Phos-tag-acrylamide, 237 μM $ZnCl_2$, 10 mM HEPES, pH 7.2, 750 mM GdnHCl). After recording for ~10 min, phosphoserine (Sigma-Aldrich) was introduced to the same compartment to a final concentration of 237 μM and another 10-min recording was performed.

Fractions (%) of phosphorylated linkers detected in A1-P-$PAZn_2$ states were calculated as:

Percentage =

$$\frac{\text{phosphorylated linkers detected in A1-P-PAZn}_2\text{ states}}{\text{phosphorylated linkers detected in A1-P-PAZn}_2\text{ states + phosphorylated linkers detected in A1-P states}}$$

If two phosphorylated linkers detected in A1-P or $PAZn_2$-bound states appeared in a single translocation event, they were analyzed individually.

**Single-channel recording**

Electrical recordings were performed with planar lipid bilayers at 23.0 ± 1.0 °C. Planar bilayers composed of 1,2-diphytanoyl-sn-glycero-3-phosphocholine (Avanti Polar Lipids) were formed by using the Müller-Montal method across a 50 μm-diameter aperture in a Teflon film (25 μm thick, Goodfellow) separating the cis and trans compartments of the recording chamber (500 μL each). Each compartment was filled with 500 μL recording buffer (10 mM HEPES, pH 7.2, 750 mM GdnHCl). Following the insertion of a single pore into the bilayer, the solution was perfused by manual pipetting to prevent further insertions. Trx-linker pentamers, Trx-linker pentamers with Phos-tag (1,3-Bis[bis(2-pyridylmethyl)amino]-2-propanol, Santa Cruz Biotechnology) dizinc complexes or Trx-linker pentamers with Phos-tag-acrylamide dizinc complexes were added to the cis compartment. For experiments in the presence of Phos-tag dizinc complexes, the phosphorylated Trx-linker pentamer was incubated with Phos-tag dizinc complex or Phos-tag-acrylamide dizinc complex at room temperature for 15 min. Then the mixture was added to the cis compartment (Trx-linker pentamers, 2.37 μM; Phos-tag or Phos-tag-acrylamide under 50 equivalents of $PZn_2$ or $PAZn_2$ conditions, 118.5 μM; $ZnCl_2$ under 50 equivalents of $PZn_2$ or $PAZn_2$ conditions, 237 μM). Ionic currents were measured using Ag/AgCl electrodes connected to a patch-clamp amplifier (Axopatch 200B, Axon Instruments). Data were low-pass Bessel filtered at 10 kHz and sampled at 50 kHz with a Digidata 1440A digitizer (Molecular Devices). Current traces were idealized by using Clampfit 10.7 (Molecular Devices). Dwell time analysis for the idealized data was performed by using the maximum interval likelihood algorithm of QuB 2.0 software (www.qub.buffalo.edu)[1,2].
